# Preventing Inappropriate Signals Pre- and Post-Ligand Perception by a Toggle-Switch Mechanism of ERECTA

**DOI:** 10.1101/2024.09.20.612365

**Authors:** Liangliang Chen, Michal Maes, Alicia M. Cochran, Julian R. Avila, Paul Derbyshire, Jan Sklenar, Kelsey M. Haas, Judit Villén, Frank L.H. Menke, Keiko U. Torii

## Abstract

Dynamic control of signaling events requires swift regulation of receptors at an active state. By focusing on Arabidopsis ERECTA (ER) receptor kinase, which perceives peptide ligands to control multiple developmental processes, we report a mechanism preventing inappropriate receptor activity. The ER C-terminal tail (ER_CT) functions as an autoinhibitory domain: its removal confers higher kinase activity and hyperactivity during inflorescence and stomatal development. ER_CT is required for the binding of a receptor kinase inhibitor, BKI1, and two U-box E3 ligases PUB30 and PUB31 that inactivate activated ER. We further identify ER_CT as a phosphodomain trans-phosphorylated by the co-receptor BAK1. The phosphorylation impacts the tail structure, likely releasing from autoinhibition. The phosphonull version enhances BKI1 association, whereas the phosphomimetic version promotes PUB30/31 association. Thus, ER_CT acts as an off-on-off toggle switch, facilitating the release of BKI1 inhibition, enabling signal activation, and swiftly turning over the receptors afterwards. Our results elucidate a mechanism fine-tuning receptor signaling via a phosphoswitch module, keeping the receptor at a low basal state and ensuring the robust yet transient activation upon ligand perception.

**Significance:** Cells perceive and process external signals through their cell-surface receptors, whose activity must be tightly maintained to prevent the spread of misinformation. How do plant cells prevent the inappropriate receptor activity? We identify a structural module within the C-terminal tail of the ERECTA (ER_CT), that inhibits the receptor pre- and post-signal activation. The ER_CT comprises of a linker and an α-helix. Before activation, ER_CT is autoinhibitory and associates with an inhibitory protein. Ligand perception triggers the transphosphorylation of ER_CT by the co-receptor, which then recruits a degradation machinery to swiftly turn over the activated receptor. Thus, we reveal an off-on-off toggle switch mechanism that finely adjusts the activity of the plant receptor, enabling the precise control over cell signaling.

## Introduction

Cell communication, both among individual cells and between cells and their environment, is fundamental to the survival and function of multicellular organisms, including higher plants. To manage this complex communication, plants have evolved an extensive superfamily of receptor-like kinases (RLKs), referred to hereafter as receptor kinases (RKs) for those with known functions. The RKs perceive and transmit external and endogenous signals, and their appropriate outputs modulate plant fitness as well as stress tolerance (1–5). A typical plant RLK contains an extracellular domain, a single transmembrane domain, and a cytoplasmic kinase domain. Based on various ectodomains, plant RLKs are classified into different subfamilies, among which those with extracellular leucine-rich repeat (LRR) domain, LRR-RLKs, comprise the largest subfamily (1, 6, 7). Primary LRR-RKs perceive signaling ligands, many of which are secreted peptides (8, 9). Well-studied examples of such LRR-RKs include the brassinosteroid (BR) receptor BRASSINOSTEROID INSENSITIVE1 (BRI1) and the pattern recognition receptor FLAGELLIN SENSING 2 (FLS2) (10, 11).

The early signaling events of receptor activation has been well documented for BRI1 and FLS2 (12). In the absence of ligands, BRI1 is maintained in a basal state by its autoinhibitory C-terminal tail and BRI1 KINASE INHIBITOR 1 (BKI1), which associates with the kinase domain of BRI1 (13, 14). Upon BR perception, BRI1 recruits its co-receptors, LRR-RKs from SOMATIC EMBRYPGENESIS RECEPTOR KINASES (SERKs) family (including BRI1-ASSOCIATED KINASE1 (BAK1)/SERK3, SERK1, and SERK4) via its extracellular LRR domain and disassociates with BKI1 to trigger the phosphorylation events within the BRI1-BAK1/SERK complex (15–17). *In vitro* and *in vivo* phosphorylation analyses have shown sequential auto- and transphosphorylation events between BRI1 and BAK1 (14, 18). Likewise, rapid heterodimerization and phosphorylation ensue between FLS2 and BAK1 upon perception of a flagellin peptide flg22 (19). Further phosphorylation analyses of BAK1/SERKs provided insight into specific phosphorylation signatures likely associated with developmental *vs.* immune signaling (20, 21). BAK1 and SERKs function as co-receptors for many other LRR-RKs and play crucial roles in plant growth and development, as well as plant immunity (17). Following the activation of receptor complexes, the signal will be attenuated through posttranslational modifications of the activated receptor kinases. The downregulation of receptor signaling is crucial to fine-tune signaling outputs. For example, two closely related plant U-box (PUB) E3 ubiquitin ligases PUB12 and PUB13 engage in the ligand-induced ubiquitination and degradation for both BRI1 and FLS2 (22, 23). However, the specific domain within these LRR-RKs that recruits PUB12/13 remains unclear.

The three Arabidopsis ER-family LRR-RKs (ER, ER-LIKE1 (ERL1) and ERL2) regulate many developmental processes, including shoot meristem homeostasis, leaf serration, inflorescence elongation, flower and ovule development, vascular differentiation, and stomatal patterning (24–32). Particularly, loss-of-function mutations in *ER* result in excessive asymmetric cell divisions during stomatal development, as well as a compact inflorescence with short pedicels (30–32). The activity of ER-family receptors is regulated by cysteine-rich secreted peptides from the EPIDERMAL PATTERNING FACTOR (EPF)/EPF-LIKE (EPFL) family: EPF2 and EPF1 are primarily perceived by ER and ERL1, respectively, to inhibit stomatal development (33–36). In contrast, EPFL9 (also known as Stomagen) promotes stomatal development as an antagonist of EPF2 through competitively binding to ER (37–39). Moreover, in developing stems, two other EPF/EPFL family peptides, EPFL4 and EPFL6, also activate ER to promote inflorescence elongation (40). It remains unknown how the ER kinase activity is regulated before the ligand perception.

Perception of EPF/EPFL peptides triggers the heterodimerization of ER with its co-receptor, BAK1/SERKs, which causes transphosphorylation between the ER and SERK family co-receptors (41). After signal activation, PUB30 and PUB31 are phosphorylated by BAK1 and subsequently ubiquitinate the ligand-activated ER, leading to its eventual degradation (42). Neither the exact sites of transphosphorylation of ERECTA by BAK1 nor how such phosphorylation events trigger the recruitment of PUB30/31 are known. Site-directed mutagenesis studies of the ER kinase domain, including those putative phosphorylation sites within the activation loop, have been performed before (20, 25). Yet, these studies failed to reveal the actual phosphorylation events signifying the receptor activation and/or inactivation.

Here, we report that the C-terminal tail domain of ER, ER_CT, functions as phosphorylation-controlled three-way off-on-off toggle switch to ensure the proper window of ER signal activation. The ER_CT possesses a characteristic a-helix and is deeply conserved through the ER orthologs and paralogs. Its removal confers receptor hyperactivity, triggers excessive inflorescence elongation, and over-inhibition of the stomatal cell lineages including increased resistance to Stomagen. The ER_CT is essential for the binding of the negative regulators, BKI1 and PUB30/31. Through mass-spectrometry, we demonstrate that BAK1 transphosphorylates ER and that the phosphonull version ER_CT (inactive state) associates with BKI1 whereas the phosphomimetic version (activated state) associates with PUB30/31. Our study elucidates the role of the ER_CT phosphomodule to limit the window of receptor activation for properly fine-tune the signal outcome and provide further insight into the modes of action of plant receptor kinases.

## Results

### ER-family receptor kinases possess a long conserved C-terminal tail

To understand the mode of ER activation, we first compared the amino acid sequence of fourteen ER orthologs from liverworts, moss, monocots, and dicots including Arabidopsis ER paralogs, ERL1 and ERL2. In addition to the leucine-rich repeats and the kinase domains, high sequence conservations were detected within the C-terminal tail region. (Fig. 1a). Notably, the last 15 amino acids of the C-terminal tail are nearly fully conserved with exception of three *Physomitrella* ERs (PpER1-3) (Fig. S1). Next, we examined the predicted Alphafold2 structures of the ER cytoplasmic domain (ER_CD) (43). The overall tail region does not adopt a discrete structure and flops out from the kinase domain. In contrast, the highly conserved last 15 amino acid C-terminal tail is predicted to form an α-Helix (Figs. 1B, S2A). This predicted α-Helix exists in all the ER paralogs in Arabidopsis and orthologs from other plant species tested (e.g., *Glycine soja* and *Oryza Sativa*) (Figs. 1, S1, S2A).

**Figure 1.**
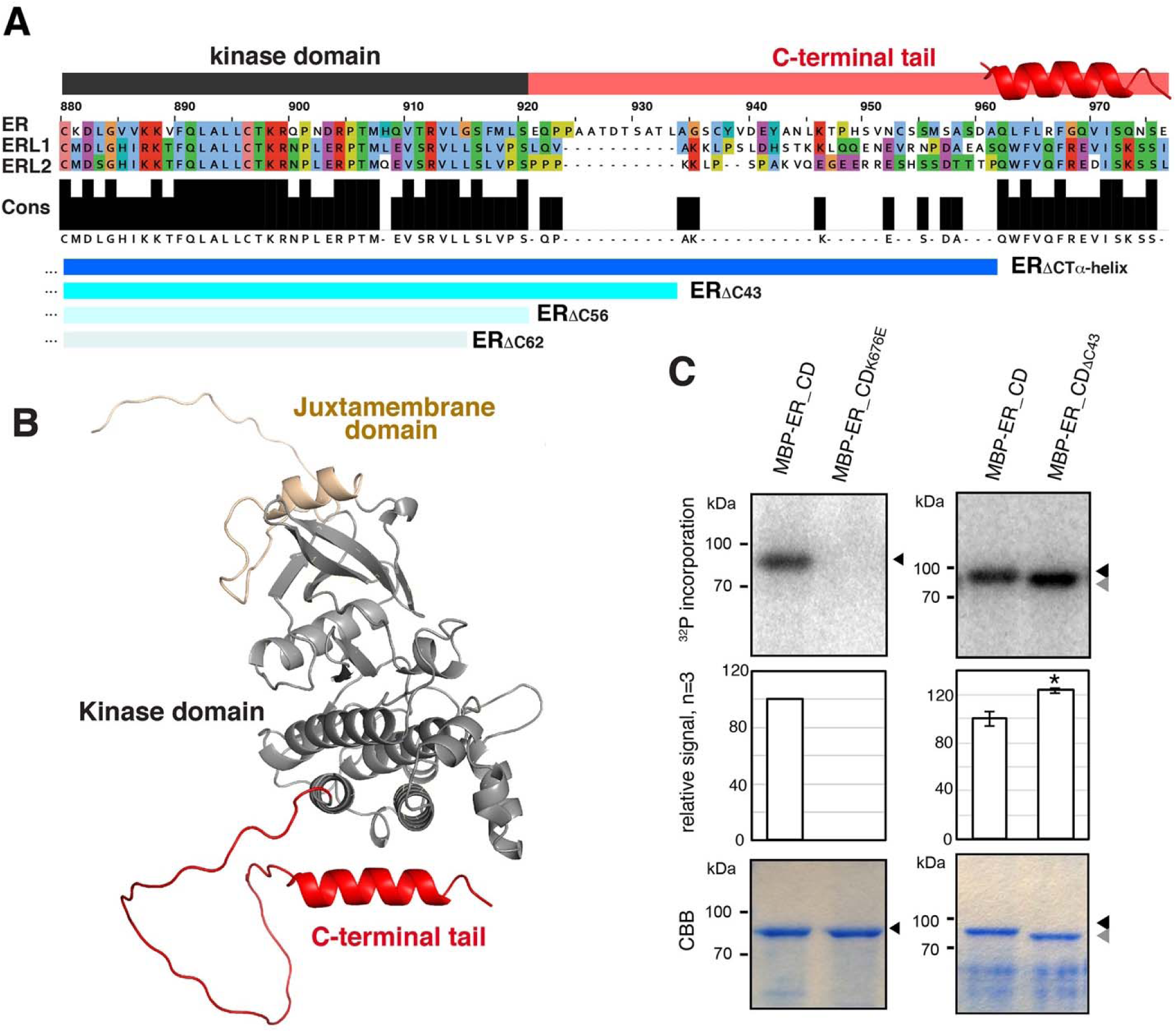
The conserved C-terminal tail in ER-family LRR-RKs inhibits the kinase activity. **(A)** Alignment of the C-terminal tail domain (ER_CT) of ER and ERLs. Conserved residues (Cons) are filled in black. Kinase domain is highlighted in dark gray, and ER_CT is in red. The location of α-Helix is superimposed. Deletions used in this study are indicated in blue: ER_ΔCTα-Helix_, ER_ΔC43_, ER_ΔC56_, and ER_ΔC62_. **(B)** Structural modeling of the cytoplasmic domain of ER (ER_CD), with the Juxtamembrane domain in sand, the Kinase domain in gray, and the C-terminal tail domain in red. **(C)** *In vitro* phosphorylation assay of ER_CD. Left panels, control ER_CD and kinase-dead version (ER_CD_K676E_). Right panels, ER_CD and ER_CD _ΔC43_. Top panels, autoradiography. Middle panels, mean and S.D. of densitometry plotted for the experiments performed 3 times. ** T-test p<0.05. Bottom panels, Coomassie Brilliant Blue staining (CBB) as a loading control.

To explore whether the sequence conservation of the C-terminal tail region extends to LRR-RK superfamily in general, we compared the tail regions of 23 representative LRR-RKs (e.g., BRI1-BRL1-BRL2-BRL3, and SERK1-SERK2-BAK1-SERK4) in Arabidopsis (Fig. S3). The alignments suggest that the sequence similarities within the C-terminal tails are relatively limited within each receptor family. Notably, the α-Helix structure is absent in C-terminal tail of other LRR-RKs tested, including BAM1, CLV1, EFR, FLS2, HSL1, HSL2, and PSKR1 (Fig. S2B). Furthermore, the tail lengths vary among these LRR-RKs, with Arabidopsis ER having the longest tail of 56 amino acids (Fig. S3). The long C-terminal tail with the terminal α-Helix may have a specific function shared within the ER subfamily.

We next asked whether the unique C-terminal tail directly affects the kinase activity of ER (Fig. 1b). For this purpose, we expressed and purified an MBP-fused full-length ER cytoplasmic domain (MBP-ER_CD), a control kinase-inactive version in which the invariable lysine residue in the ATP binding domain is substituted to a glutamate (MBP-ER_CD_K676E_), and MBP-ER_CD_ΔC43_, in which the C-terminal 43 amino acids are removed. Subsequently, *in vitro* kinase assays were performed. MBP-ER_CD exhibited autophosphorylation, while MBP-ER_CD_K676E_ showed no autophosphorylation (Fig. 1C). Notably, MBP-ER_CD_ΔC43_ showed slight yet reproducible increase of autophosphorylation (Fig. 1C). The results suggest that the C-terminal tail negatively regulates ER kinase activity.

### The C-terminal tail negatively regulates ER function

The ER C-terminal tail (ER_CT) was proposed to be dispensable for the ER function (25). However, the increased kinase activity in the absence of ER_CT suggests its potential regulatory function (Fig. 1C). To test this hypothesis, we first introduced three truncated versions of *ER* driven by its own promoter into *er* null allele, *er-105*: *ER*_Δ*CT*α_*_-Helix_, ER_ΔC56_*, and *ER_ΔC62_* lacking the C-terminal α-helix, the entire tail, and the tail plus the six last amino acids within the kinase domain, respectively (Fig. 1A). *ER*_Δ*CT*α*-Helix*_ and *ER_ΔC56_* not only rescued *er* inflorescence phenotype, but also increased their heights to approximately ∼120% of control wild-type plants (Fig. S4A, B), suggesting that removal of the C-terminal tail confers hyperactivity of ER. On the other hand, *ER_ΔC62_* failed to rescue the compact *er* inflorescence phenotype (Figs. S4A, B), consistent with the importance of ER kinase activity for its function (30).

We next examined the effects of these ER_CT deletions on stomatal development (Fig. 2). In *er*, clusters of small stomatal-lineage ground cells (SLGCs) are produced due to excessive entry divisions of stomatal-lineage cells (32) (Fig. S4C). The stomatal densities (number of stomata/unit area) of *ER*_Δ*CT*α*-Helix*_ and *ER_ΔC56_,* but not *ER_ΔC62_*, are lower than wild type (Fig. S4D), suggesting that removal of the ER_CT overly represses stomatal differentiation. It is known that ER signaling pathway leads to downregulation of the transcription factor SPEECHLESS (SPCH), which initiates stomatal cell lineages (44, 45). To further characterize the effects of *ER_ΔC_* on stomatal development at a molecular level, we further analyzed the expression levels of *SPCH* and known direct SPCH targets, *EPF2*, *TMM*, and *POLAR* (46, 47) (Fig. S4E-G). Both *ER*_Δ*CT*α*-*_ *_Helix_* and *ER_ΔC56_* reduced the transcript levels of these genes, some with statistical significance. Thus, removal of ER_CT, either the entire domain or the α-helix alone, is also sufficient to confer hyperactivity of ER in stomatal development.

**Figure 2.**
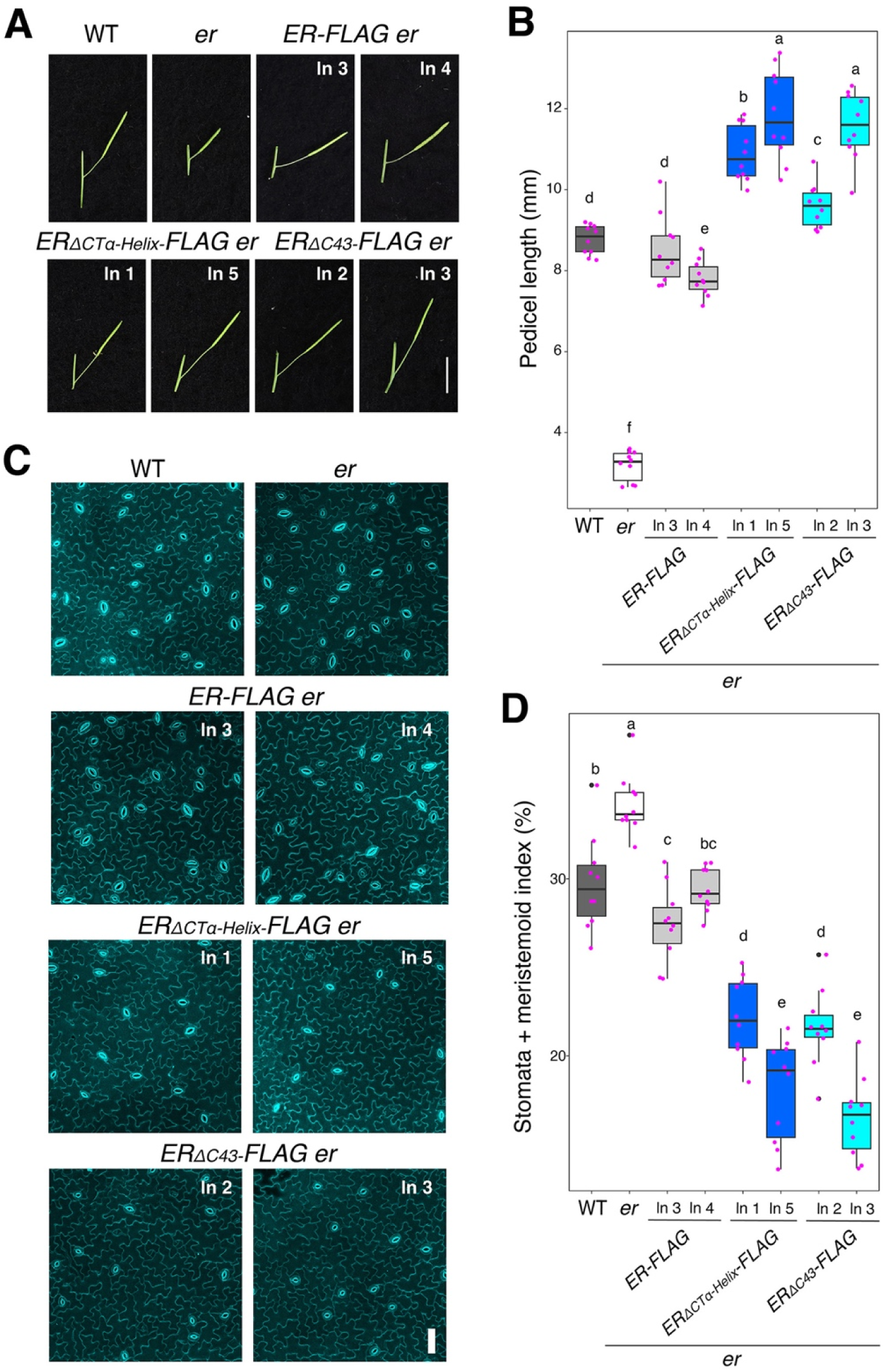
Removal of the ER C-terminal tail domain leads to hyperactivity in promoting pedicel elongation and inhibiting stomatal development. **(A)** Representative pedicels and mature siliques of WT, *er-105 (er)*, *ER-FLAG er*, *ER*_Δ*CT*α_*_-Helix_-FLAG er*, and *ER*_Δ*C43*_*-FLAG er* plants. For each transgenic constructs, two representative lines were subjected to analysis. Images were taken under the same magnification. Scale bar, 1 cm. **(B)** Morphometric analysis of pedicel length from each genotype. Six-week-old mature pedicels (n = 10) were measured. One-way ANOVA followed by Tukey’s HSD test was performed and classified their phenotypes into categories (e.g. a, b, c). **(C)** Representative confocal microscopy of cotyledon abaxial epidermal at 7 day-post-germination (DPG7) from wild type (WT), *er-105* (*er*), *ER-FLAG er*, *ER*_Δ*CT*α_*_-Helix_-FLAG er*, and *ER*_Δ*C43*_*-FLAG er.* The same representative transgenic lines were observed. Images were taken under the same magnification. Scale bar = 50 µm. **(D)** Quantitative analysis. Stomata + meristemoid index (number of stomata and meristemoid per 100 epidermal cells) of the cotyledon abaxial epidermis from 7-day-old seedlings of respective genotypes (n = 10). One-way ANOVA followed by Tukey’s HSD test was performed and classified their phenotypes into categories (e.g. a, b, c).

Now we know that ER_CT negatively regulates the biological function of ER, we subsequently generated *er* plants expressing the epitope-tagged versions of the truncated ER_CT for further biochemical analyses. For this purpose, we focused on the deletions of conserved α-helix domain, *ERpro::ER*_Δ*CT*α_*_-Helix_-FLAG,* and that enhances the *in vitro* kinase activity, *ERpro::ER*_Δ*C43*_*-FLAG*. Again, both *ERpro::ER*_Δ_*_CT_*_α_*_-Helix_-FLAG*, and *ERpro::ER*_Δ*C43*_*-FLAG* conferred over-complementation with excessive pedicel elongation (Fig. 2A, B) and reduced numbers of stomatal and meristemoid (stomata + meristemoid index = (number of stomata and meristemoid)/number of stomata + meristemoid + all other epidermal cells) x100) compared to the control *ER-FLAG* lines (Fig. 2C, D). The transcript levels of the transgenes are comparable among the transgenic lines expressing *ER-FLAG*, *ER*_Δ*CT*α_*_-Helix_-FLAG,* and *ER*_Δ*C43*_*-FLAG* (Fig. S5A), indicating that the observed excessive phenotypic rescues are not attributable to the transgene overexpression. Notably, the protein level of ER deletion versions is higher than the wild-type version, indicating the C-terminal tail regulates ER protein level (Fig. S5B). Based on these findings, we conclude that the ER_CT prevents the excessive activation of ER signaling and that C-terminally truncated versions of ER, with or without the epitope tag, confer hyperactivity in the multiple contexts of ER-mediated developmental processes.

### The C-terminal tail recruits the inhibitory protein of ER

Several possible mechanisms may explain the negative regulation of ER function by its C-terminal tail. It has been reported that BRI1 KINASE INHIBITOR1 (BKI1) inhibits ER kinase activity and regulates plant architecture together with ER (48). Therefore, we asked whether ER_CT mediates the interaction between BKI1 and ER. We first quantitatively characterized the kinetics of protein-protein interactions between BKI1 and ER_CD variants via biolayer interferometry (BLI) (see Supplementary Methods). The BLI assay showed that BKI1 binds with ER_CD with equilibrium dissociation constant (Kd) values at 3.7± 0.7 μM (Fig. 3A). The interaction of BKI1 with ER_CD_ΔCTα-Helix_ was reduced roughly by 10-fold (Kd = 47.7 ± 34.1 μM), and no interaction was detected for BKI1 and ER_CD_ΔC43_ (Fig. 3A). These results suggest that the ER_CT is necessary for the association with BKI1.

**Figure 3.**
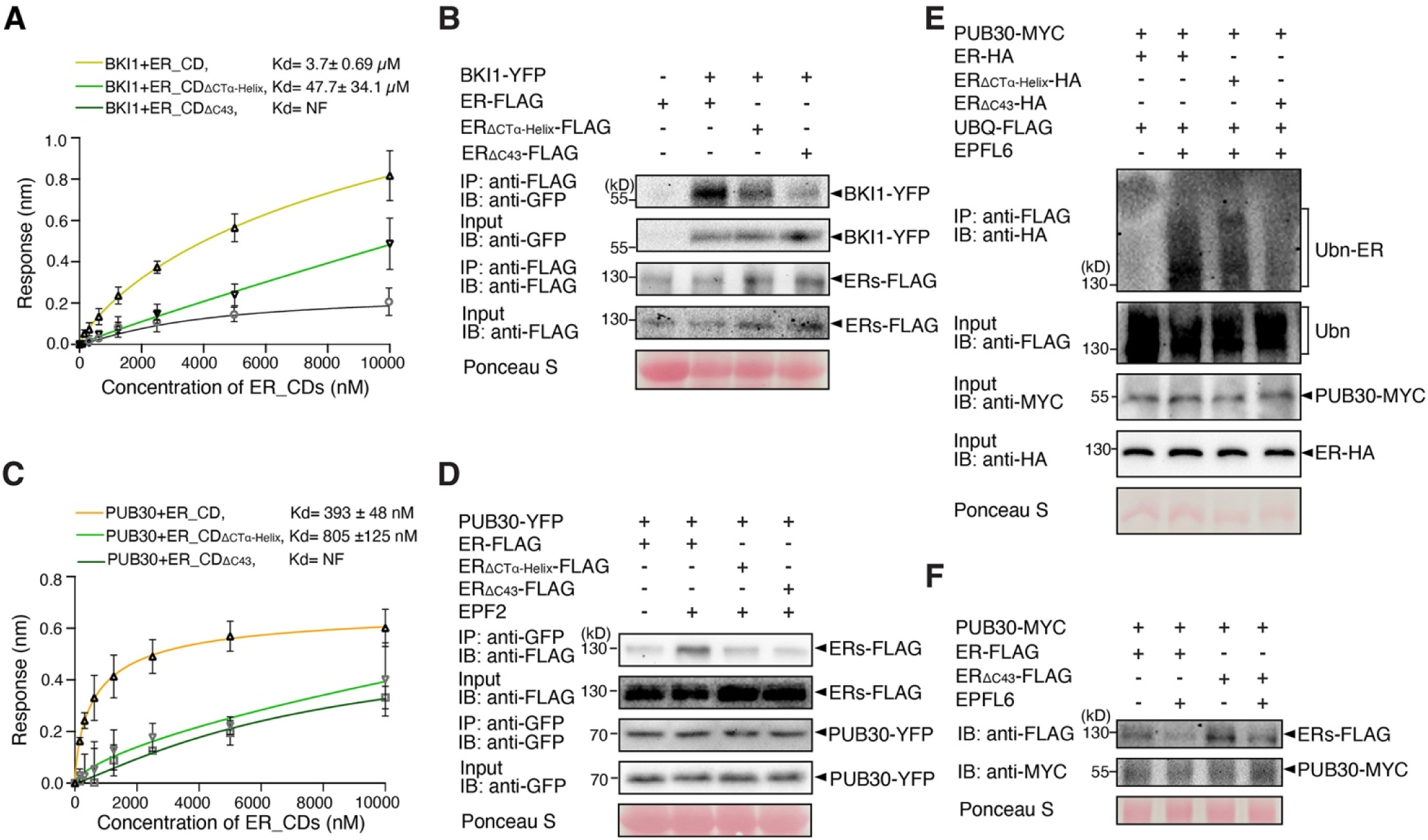
The C-terminal tail is essential for the binding of ER with BKI1 and PUB30. **(A)** Quantitative analysis of interactions between BKI1 and ER_CD variants (ER_CD, ER_CD_ΔCTα-Helix_, and ER_CD_Δ43_) using BLI. *In vitro* binding response curves for recombinantly purified GST-BKI1 and MBP-ER_CD variants at seven different concentrations (156.25, 312.5, 625, 1,250, 2,500, 5,000, and 10,000 nM) are shown. Kd values are indicated. Data are representative of three independent experiments. **(B)** Deletion of C-terminal tail decreases the association of ER with BKI1 *in vivo*. Proteins from double transgenic lines carrying *ERpro::BKI1-YFP ERpro::ER-FLAG*, *ERpro::BKI1-YFP ERpro::ER*_Δ*CT*α_*_-Helix_-FLAG*, and *ERpro::BKI1-YFP ERpro::ER*_Δ*C43*_*-FLAG* were immunoprecipitated with anti-FLAG beads (IP). The immunoblots (IB) were probed with anti-FLAG and anti-GFP antibodies, respectively. **(C)** Quantitative analysis of interactions between PUB30 and ER_CD variants (ER_CD, ER_CD_ΔCTα-Helix,_ and ER_CD_ΔC43_) using BLI. *In vitro* binding response curves for recombinantly purified GST-PUB30 and MBP-ER_CD variants at seven different concentrations (156.25, 312.5, 625, 1,250, 2,500, 5,000, and 10,000 nM) are shown. Kd values are indicated. Data are representative of three independent experiments. **(D)** Deletion of C-terminal tail decreases the association of ER with PUB30 *in vivo*. Proteins from *PUB30pro::PUB30-YFP; ERpro::ER-FLAG*, *PUB30pro::PUB30-YFP; ERpro:: ER*_Δ*CT*α_*_-Helix_-FLAG*, and *PUB30pro::PUB30-YFP; ERpro:: ER*_Δ*C43*_*-FLAG* plants were immunoprecipitated with anti-GFP beads (IP), and the immunoblots (IB) were probed with anti-GFP and anti-FLAG antibodies, respectively. **(E)** Deletion of C-terminal tail decreases the ubiquitination of ER by PUB30 *in vivo.* Arabidopsis protoplasts were co-transfected with PUB30-MYC, FLAG-UBQ, together with ER-HA, ER_ΔCTα-Helix_-HA, and ER_ΔC43_-HA. Five micromolar EPFL6 was used for treatment for 1 hr. After immunoprecipitation using anti-FLAG beads, the ubiquitinated ER variants were probed with anti-HA antibody. The total ubiquitinated proteins were probed by anti-FLAG antibody and PUB30 proteins were probed by anti-MYC antibody. The inputs of ER were probed with anti-HA antibody. **(F)** Representative EPFL6 treatment destabilizes ER variants in Arabidopsis protoplasts co-expressing PUB30-MYC. Protoplasts expressing the indicated proteins were treated with 50 μM CHX and 5 μM EPFL6 for 3 hr. before total protein was examined with immunoblot. The experiment was repeated independently two times with similar results.

Next, to examine the *in vivo* association of BKI1 and ER variants in Arabidopsis, we performed co-immunoprecipitation (Co-IP) analyses using transgenic plants carrying epitope-tagged ER variants (*ERpro::ER-FLAG*, *ERpro::ER*_Δ*CT*α_*_-Helix_-FLAG*, and *ERpro::ER*_Δ*C43*_*-FLAG*) and BKI1 (*ERpro::BKI1-YFP*). Much less BKI1-YFP signal was detected in the immunoprecipitated ER_ΔCTα-Helix_-FLAG or ER_ΔC43_-FLAG than ER-FLAG complexes (Fig. 3B), indicating that ER_CT is required for the *in vivo* binding of ER with BKI1. It has been reported that EPF/EPFL perception by ER triggers the formation and activation of the receptor complex (41, 42). To test a hypothesis that receptor activation inhibits the interaction of BKI1 and ER, we furthermore treated the seedlings with bioactive, mature recombinant EPF2 (MEPF2) peptides. Indeed, association of ER with BKI1 decreased upon MEPF2 application (Fig. S6). Combined, our results demonstrate that the C-terminal tail of ER is required for its interaction with BKI1 and that the ER signal activation triggers the dissociation of BKI1.

### The C-terminal tail negatively regulates ER protein abundance

Our previous work illustrated that PUB30 and PUB31 downregulate the accumulation of ligand stimulated ER proteins via ubiquitination. We then asked whether ER_CT is required to recruit these E3 ligases. We found that ER_CD tightly binds with PUB30 with Kd values at 393 ± 48 nM in the BLI assays. Removal of the α-Helix reduced the association with PUB30, increasing the Kd value by a factor of three (805 ± 125 nM). Notably, ER_CD_ΔC43_ did not show any detectable binding with PUB30 (Fig. 3C). Like PUB30, PUB31 also exhibits weaker interaction with ER_CD C-terminal deletion version compared with ER_CD (Fig. S7A).

Next, to test their *in vivo* protein-protein interactions, we performed Co-IP analyses using transgenic plants expressing FLAG-tagged ER_CD variants and PUB30 (*PUB30pro::PUB30-YFP*), as well as protoplasts co-expressing epitope-tagged ER variants (ER-HA, ER_ΔCTα-Helix_-HA, and ER_ΔC43_-HA) and PUB31 (PUB31-MYC) (see SI Materials and Methods). Compared to the ER, much less amounts of ER_ΔCTα-Helix_ or ER_ΔC43_ proteins were detected when subjected to Co-IP with PUB30-YFP or PUB31-MYC (Figs. 3D, S7B), indicating that, in addition to the binding with BKI1, the ER C-terminal tail is also required for the binding with PUB30/31.

To address whether ER_CT is required for turn-over of ligand-activated ER, we subsequently performed an *in vivo* ubiquitination assay using Arabidopsis protoplasts co-expressing epitope-tagged ER (ER-HA), PUB30 or PUB31 (PUB30-MYC or PUB31-MYC) and ubiquitin (FLAG-UBQ) (see SI Materials and Methods). Laddering bands with high molecular mass proteins are detected after immunoprecipitation, indicative of the ubiquitination of ER *in vivo* (Figs. 3E, S7C). Strikingly, the deletion of ER_CT, most evident in ER_ΔC43,_ diminishes the polyubiquitination of ER by PUB30 and PUB31 (Figs. 3E, S7C), suggesting that ER_CT is required for the ligand-stimulated ER ubiquitination. Finally, we tested the effects of ER_CT on the degradation of ER protein. For this purpose, we co-expressed FLAG-tagged ER variants and PUB30 in Arabidopsis protoplast and performed cotreatment with bioactive EPFL6 peptide (MEPFL6) in the presence of cycloheximide (*de novo* protein synthesis inhibitor). MEPFL6 treatment substantially decreased the accumulation of ER protein but not as much of the ER_ΔC43_ protein (Fig. 3F, S7B). Based on these findings, we conclude that ER_CT is required for the interaction with PUB30/31, the ubiquitination by PUB30/31, and the turn-over of ligand stimulated ER proteins.

### The ER C-terminal tail is an *in vivo* phosphodomain

In animal receptor kinases, phosphorylation of C-terminal tail plays a role in signal transduction (49). *In vivo* and *in vitro* phosphorylation of the C-terminal tail has been reported for BRI1 and BAK1 (14, 18, 20, 50, 51). The long, conserved C-terminal tail domain of ER orthologs possesses multiple serines and threonines that could serve as phospho-acceptor residues, many of which are highly conserved (Fig. S1). In contrast, neither BRI1 nor BAK1 has such extended serine/threonine-rich tails (Fig. S3). This leads us to investigate whether the ER_CT is subjected to phosphorylation and, if so, what is their functional significance.

To identify *in vivo* ER phosphosites, we performed affinity purification of Arabidopsis ER protein using transgenic Arabidopsis *er* seedlings fully complemented by functional *ER* fused with YFP driven by the endogenous promoter (*ERpro::ER-YFP*). Immunoprecipitated Arabidopsis ER-YFP protein was subjected to LC-MS/MS analysis. Eighteen experiments were performed using three different mass spectrometry experimental settings (see SI Materials and Methods; Fig. S8 for total coverage). We detected phosphosites at Thr947, Ser955, Ser972 as well as additional phosphosites Ser950, Ser954, Ser957 with less confidence in localization (Figs. 4, S9, Table S1). All residues are removed in ER_ΔC43_. Among them, Ser955, Ser957, and Ser972 are highly conserved among the ER orthologs and paralogs (Fig. S1), and Thr947 and Ser950 share conservation among the ER orthologs in Angiosperms and *Brassicaceae*, respectively (Fig. S1). Thus, the highly conserved ER_CT defines an *in vivo* phosphodomain. Additional phosphosites were identified within the activation loop of ER kinase domain, Ser801 and Tyr808, which are indispensable for the ER function (25), as well as Ser680 and Ser766. The recoveries of these phosphorylated peptides were low, however, implying that the ER kinase domain itself may not be pre-dominantly phosphorylated *in vivo*.

**Figure 4.**
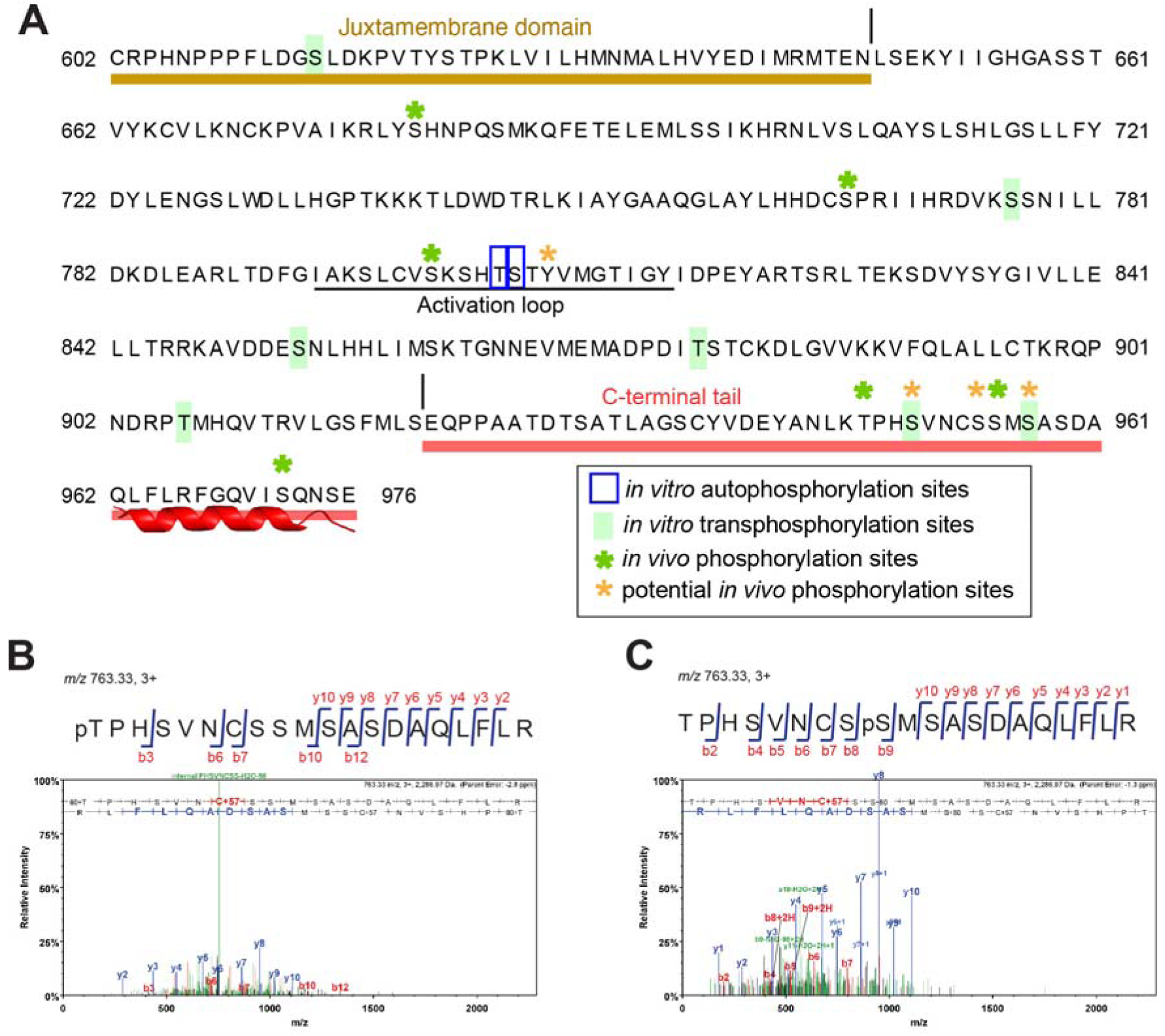
The C-terminal tail domain of ER is phosphorylated. **(A)** Phosphorylation sites in the C-terminal tail domain of ER. *In vitro* auto- and trans-phosphorylation sites identified by LC-MS/MS analysis are marked in blue and lime green, respectively. *In vivo* localized phosphorylation sites of ER found in this study are marked with green asterisks. Potential *in vivo* phosphorylation sites of ER are marked with orange asterisks. Juxtamembrane and C-terminal tail are highlighted in sand and red, respectively. Underline, the Activation Loop. **(B, C)** MS/MS spectra for selected *in vivo* phosphorylation sites of ER: Thr947 (**B**) and Ser955 (**C**).

### The ER C-terminal tail is transphosphorylated by BAK1

Our work revealed that removal of ER_CT confers hyperactivity of ER (Figs. 1-3). Furthermore, we identified multiple *in vivo* phosphosites on the C-terminal tail (Fig. 4, Table S1). To delve into the underlying mechanism of phosphoregulation, we first analyzed ER autophosphorylation using MBP-ER_CD (Fig. 1C) (52) by mass-spectrometry analysis. In contrast to the *in vivo* data, minimal *in vitro* phosphorylation was detected only within the ER activation loop (Thr805, Ser806; Figs. 4A, Table S1), indicating that ER_CD is not autophosphorylated. As expected, no phosphopeptides were detected in the control kinase-dead ER (MBP-ER_CD_K676E_, Table S1).

Although it has been shown by autoradiography that BAK1 can transphosphorylate ER (41), the exact phosphosites are unknown. Our mass spectrometry analysis after co-incubation of MBP-ER_CD with a recombinant BAK1 cytoplasmic domain (MBP-BAK1_CD), identified unambiguous phosphorylation sites at Ser950 and Ser957, and low localization score at Ser972 within the ER_CT (Figs. 4, S9, Table S1). The high-confidence C-terminal tail phosphorylation occurred even when the kinase-dead ER protein (MBP-ER_CD_K676E_) was co-incubated with BAK1, thereby demonstrating that the ER_CT is transphosphorylated by BAK1. Importantly, all of these *in vitro* transphosphorylation sites correspond to the *in vivo* ER phosphorylation sites (Fig. 4, Table S1). In addition to the ER phosphosites, we also detected strong autophosphorylation of BAK1 *in vitro* (Table S2) within the activation loop, which is required for kinase activation (21), as well as in the juxtamembrane and C-terminal tail domains (Table S2). Importantly, *in vitro* BAK1 phosphosites detected in our study include all of the previously reported *in vivo* BAK1 phosphosites within the kinase domain (Table S2) (14, 18, 20, 21).

### Phosphorylation of the C-terminal tail as a switch for the binding with BKI1 to PUB30/31

Our biochemical studies revealed that ER_CT negatively regulates the kinase activity and protein abundance of ER through recruiting BKI1 and PUB30/31 before and after the activation of ER, respectively (Figs. 2, 3). To address whether phosphorylation status of the C-terminal tail affect the interaction of ER with BKI1 and PUB30/31, we performed site-directed mutagenesis of the eight phosphosites within ER_CT, including six experimentally detected sites and two potential sites (Ser959 and Ser975), for a series of *in vitro* and *in vivo* assays. The *in vitro* BLI assays showed that the phosphomimetic version ER_CD_T/S8E_ exhibits weaker interaction with BKI1 than the wild-type ER_CD or the phosphonull version ER_CD_T/S8A_ (Fig. 5A). Next, we performed *in vivo* Co-IP experiments using transgenic plants carrying epitope tagged-ER, *ERpro::ER-FLAG,* phosphonull version *ERpro::ER_T/S8A_-FLAG*, and phosphomimetic version *ERpro::ER_T/S8E_-FLAG* with BKI1 (*ERpro::BKI1-YFP*). The association of BKI1-YFP with ER_T/S8E_-FLAG was markedly reduced compared with the wild-type version (Fig. 5B). By contrast, the phosphonull mutant ER_T/S8A_-FLAG exhibited stronger interaction with BKI1 than the wild-type ER-FLAG (Fig. 5B). These results suggest that phosphorylation within ER_CT abolishes the binding of BKI1 and ER. Intriguingly, the phosphonull version, ER_CD_T/S8A_, exhibits much weaker interaction with PUB30 than the wild-type version ER_CD or the phosphomimetic version ER_CD_T/S8E_ (Fig. 5C). Further *in vivo* co-immunoprecipitation assays from Arabidopsis seedlings demonstrated both PUB30 and PUB31 show weak interaction with the phosphonull version of ER (ER_T/S8A_-FLAG) (Fig. 5D). In contrast, PUB30 and PUB31 show stronger interaction with the phosphomimetic version of ER (ER_T/S8E_-FLAG) than the wild-type ER-FLAG, thereby confirming that phosphorylation of the C-terminal tail intensifies the association with its E3 ligases (Fig. 5D). Collectively, these findings suggest that the phosphorylation of the ER C-terminal tail leads to the eviction of BKI1 and recruitment of PUB30/31.

**Figure 5.**
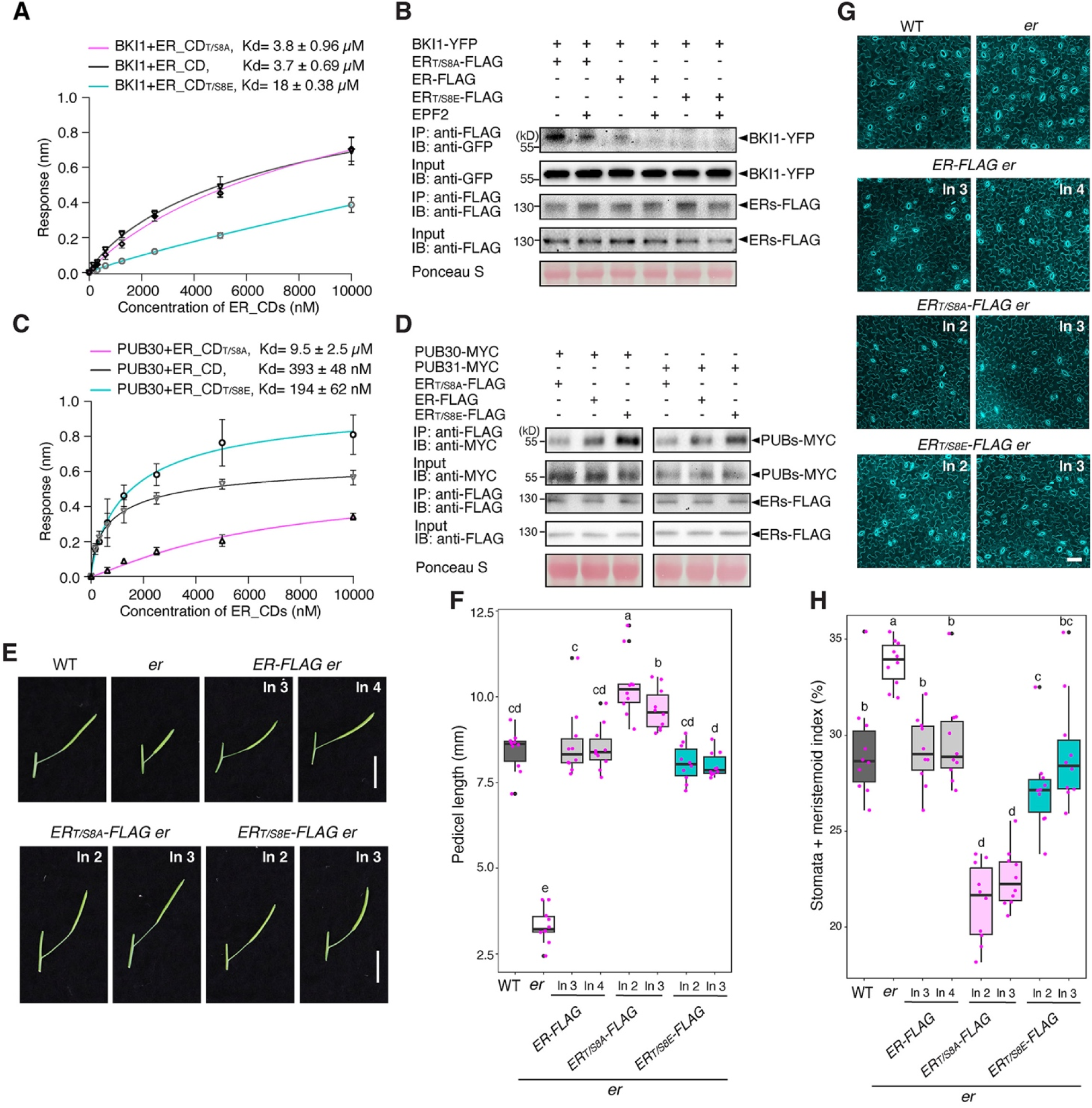
Phosphorylation of the ER C-terminal tail evicts BKI1 and recruits PUB30/31. **(A)** Quantitative analysis of interactions between BKI1 and ER_CD variants (ER_CD, ER_CD_T/S8A_, and ER_CD_T/S8E_) using BLI. *In vitro* binding response curves for recombinantly purified GST-BKI1 and MBP-ER_CD variants at seven different concentrations (156.25, 312.5, 625, 1,250, 2,500, 5,000, and 10,000 nM) are shown. Kd values are indicated. Data are representative of three independent experiments. **(B)** Phosphorylation of C-terminal tail decreases the association of ER with BKI1 *in vivo*. Proteins from *ERpro::BKI1-YFP; Col-0, ERpro::BKI1-YFP; ERpro::ER-FLAG*, *ERpro::BKI1-YFP; ERpro::ER_T/S8A_-FLAG*, and *ERpro::BKI1-YFP; ERpro::ER_T/S8E_-FLAG* plants were immunoprecipitated with anti-FLAG beads (IP), and the immunoblots (IB) were probed with anti-FLAG and anti-GFP antibodies, respectively. **(C)** Quantitative analysis of interactions between PUB30 and ER_CD variants (ER_CD, ER_CD_T/S8A_, and ER_CD_T/S8E_) using BLI. *In vitro* binding response curves for recombinantly purified GST-PUB30 and MBP-ER_CD variants at seven different concentrations (156.25, 312.5, 625, 1,250, 2,500, 5,000, and 10,000 nM) are shown. Kd values are indicated. Data are representative of three independent experiments. **(D)** Phosphorylation of C-terminal tail promotes the association of ER with PUB30 and PUB31 *in vivo*. Arabidopsis protoplasts were co-transfected with PUB30-MYC or PUB31-MYC, together with ER-FLAG, ER_T/S8A_-FLAG, and ER_T/S8E_-FLAG. Five micromolar EPFL6 was used for treatment for 1 h. After immunoprecipitation using anti-FLAG beads, the immunoblots (IB) were probed with anti-FLAG and anti-MYC antibodies, respectively. **(E)** Representative pedicels and mature siliques of WT, *er*, *ER-FLAG er*, *ER_T/S8A_-FLAG er*, and *ER_T/S8E_-FLAG er* plants. Images were taken under the same magnification. Scale bar, 1 cm. **(F)** Morphometric analysis of pedicel length from each genotype. Six-week-old mature pedicels (n = 10) were measured. One-way ANOVA followed by Tukey’s HSD test was performed and classified their phenotypes into categories (a, b, c, and d). **(G)** Confocal microscopy of 7-day-old abaxial cotyledon epidermis of WT, *er*, *ER-FLAG er*, *ER_T/S8A_-FLAG er*, and *ER_T/S8E_-FLAG er* plants. Images were taken under the same magnification. Scale bar, 50 μm. **(H)** Quantitative analysis. Stomata + meristemoid index of the cotyledon abaxial epidermis from 7-day-old seedlings of respective genotypes (n = 10). One-way ANOVA followed by Tukey’s HSD test was performed and classified their phenotypes into categories (a, b, c, and d).

### Phosphorylation of the C-terminal tail activates and attenuates the ER signaling

To further evaluate the contribution of the C-terminal tail phosphorylation on biological functions of ER, we introduced the C-terminal tail phosphomimetic and phosphonull versions of ER driven by its native promoter into *er* null mutant plants. Transgenic *er* plants expressing the phosphomimetic version *ERpro::ER_T/S8E_-FLAG* rescued both pedicel growth and stomatal phenotype of *er* in a similar degree to the complementation lines expressing the wild-type version *ERpro::ER-FLAG* (Fig. 5E, F). Remarkably, transgenic plants expressing the phosphonull version *ERpro::ER_T/S8A_-FLAG* overly rescued the *er* mutant phenotypes, both in the context of pedicel growth and stomatal development (Fig. 5E, F). Thus, having a C-terminal tail that cannot be phosphorylated mimics the hyperactivity of ER lacking the C-terminal tail (e.g. ER_ΔCTα-Helix_ and ER_ΔC43_) (see Figs. 1-2, S4). To exclude the possibility that the various degrees of transcript levels lead to the differences in phenotypic rescues by the phosphonull *ER_T/S8A_*, we examined both transcriptional levels in these transgenic lines. The transcriptional levels of phosphomimetic, wild-type, and phosphonull versions of *ER* are comparable (Fig. S10A). Notably, the protein level of phosphonull version is higher than the wild-type version, while that of the phosphomimetic version is lower (Fig. S10B). To confirm the effects of phosphorylation in ER_CT on the stability of ER protein, we co-expressed FLAG-tagged ER phosphovariants and PUB30 in Arabidopsis protoplast and performed co-treatment with bioactive EPFL6 peptide in the presence of cycloheximide. MEPFL6 application substantially decreased the accumulation of wild-type ER protein but not as much of the ER_T/S8A_ protein (Fig. S11A). Conversely, the level of ER_T/S8E_ protein was markedly lower compared to the wild-type ER protein, in both the absence and presence of ligand perception (Fig. S11B). Taken together, our results highlight that phosphorylation of ER_CT underscores the biological functions of ER.

### Phosphorylation within the ER C-terminal tail impacts its structure

The AlphaFold prediction of ER_CT adopts a flexible structure, ending with an α-Helix (Figs. 1, S3). The Ser972 residue, one of the *in vivo* and *in vitro* transphosphorylation sites, is located within the ER_CTα-Helix. This serine residue is highly conserved among the ER orthologs and paralogs (Fig. S1), and its phosphorylation is predicted to affect the flexibility of the α-Helix (Fig. 6A). We postulated that the phosphorylation within the ER_CT may impact its secondary structure. To explore this possibility experimentally, we employed circular dichroism (CD) spectroscopy to analyze the secondary structures of a synthetic peptide (ER_CTα-Helix) and its phosphorylated form at the Ser972 residue (ER_CTα-HelixS972p) (Fig. 6B). The CD spectra confirmed the dominance of the predicted α-Helix in the last 15 aa of the ER_CT. Further secondary structure analysis indicates that the Ser972 phosphorylation reduces the Helix content from 76.2 % to 69.7 % (Fig. 6B), thus verifying the modeling prediction. Based on these findings, we propose that phosphorylation of ER_CT by BAK1, triggered by the EPF/EPFL ligand perception and subsequent ER-BAK1 heterodimerization, induces a structural change that alleviates ER autoinhibition and promotes the dissociation of BKI1.

**Figure 6.**
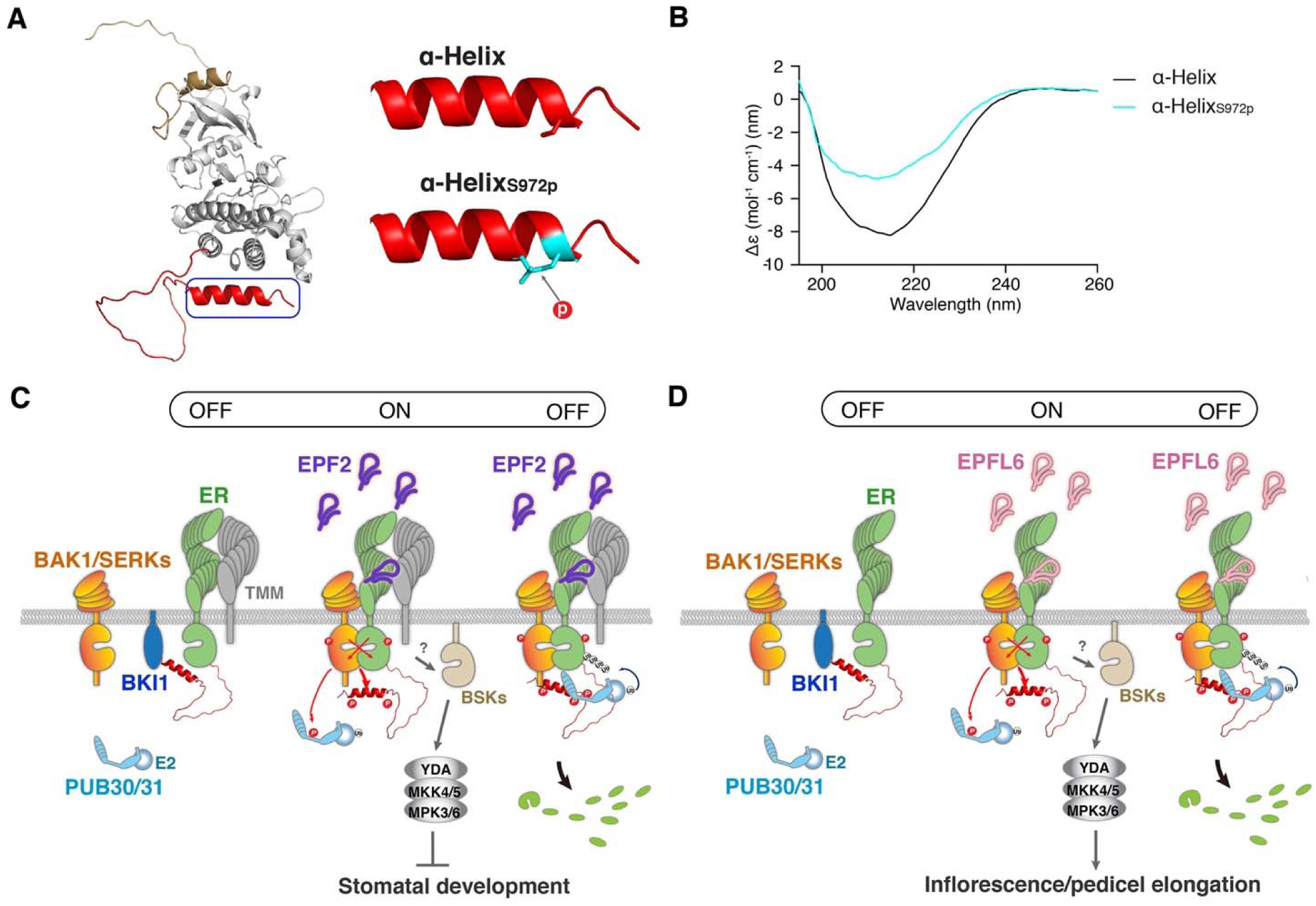
Mechanism preventing inappropriate signals pre- and post-activation of ER. **(A)** Structural modeling of ER_CD (left). Structural modeling of ER_CTα-Helix domain (right, top), with the Ser972 residue shown as sticks in red, and ER_CTα-Helix_S972p_ (right, bottom), with the phosphorylated Ser972 residue shown as sticks in cyan. **(B)** CD spectra of ER_CTα-Helix (black) and ER_CTα-Helix_S972p_ (cyan). **(C, D)** Regulation of stomatal development (**C**) and inflorescence/pedicel elongation (**D**) by ER_CT. (Left) BKI1 (blue) associates with ER (green) in the absence of ligand. (Middle) Upon perception of EPF2 (**C**, violet) or EPFL6 (**D**, pink), ER becomes activated. Both ER kinase domain and C-terminal tail are phosphorylated by its co-receptor BAK1/SERKs (orange) during the transphosphorylation events. The phosphorylation of C-terminal tail evicts BKI1. Activated BAK1/SERKs also phosphorylate PUB30/31 (cyan). The activated ER-BAK1/SERKs receptor complex transduces signals most likely via BSK (sand) before the activation of a MAPK cascade and subsequent inhibition of stomatal development (**C**) or promotion of inflorescence/pedicel elongation (**D**). TMM (gray) biases the signal activation for stomatal development (**C**). The identity of E2 ligase (cyan ball) is unclear.

## Discussion

Based on our findings, we propose a regulatory mechanism of the cell-surface receptor as an off-on-off toggle switch, enabling the swift transition of signaling from inhibition, activation, to attenuation in response to ligand perception (Fig. 6C). A series of the biochemical analyses illustrates that the C-terminal tail of ER possesses a dual role in recruiting both the inhibitory protein BKI1 and the E3 ligases, PUB30/31, in the basal and activated state, respectively. The full activation of ER-BAK1 signaling via transphosphorylation of the ER C-terminal tail likely evokes a structural change within its α-helix (Fig. 6A, B). This event involves the relief of both intramolecular inhibition and intermolecular interactions with BKI1 (Figs. 5, 6). Thereafter, the phosphorylation events trigger association of ER with E3 ligases, which eventually leads to the turn-over of ER (Fig 5 and Fig S10). Our work sheds light not only on the mechanism of receptor autoinhibition but also that of receptor activation and attenuation orchestrated by the multi-layered phosphoregulation of a receptor complex.

It is widely accepted that phosphorylation events in the C-terminal tails are highly relevant to the activity of receptor-kinase signal transduction (13, 49, 53). However, the exact modes of actions appear quite distinct, even among the plant LRR-RKs. For instance, we have shown that ER C-terminal tail can be transphosphorylation by BAK1 (Table S1, 2), suggesting that it is part of a signal transduction process after BAK1 associates with ER upon ligand perception. The BAK1-mediated phosphorylation of ER C-terminal tail further dissociates BKI1. In contrast, a well-known LRR-RK, BRI1, exhibits strong intermolecular transphosphorylation of its own C-terminus *in vitro*, suggesting that its C-terminal tail phosphorylation signifies the initial activation step of the BRI1 homodimers (14). Subsequent BAK1-mediated BRI1 transphosphorylations then fully activates the BR signaling (18). In fact, BAK1 *per se* was not able to phosphorylate BRI1 C-terminal tail, but BAK1 could enhance BRI1 kinase activity (18). Thus, although both BRI1 and ER share BAK1/SERKs as heterodimeric partners (15, 16, 41), the exact mechanism of receptor activation and subsequent signaling may be distinct. This idea echoes a recent finding that different BAK1 phosphocodes specify BRI1 *vs*. FLS2 LRR-RK signaling (20). More recently, BAK1-mediated phosphorylation of an LRR-RK, BAK-TO-LIFE2 (BTL2), was shown to function as a phosphoswitch to prevent autoimmunity (54). This phosphorylation occurs within the juxtamembrane domain (but not the C-terminus) of BTL2, therefore highlighting the diverse mechanisms of LRR-RK phosphoregulation.

Our work further revealed that the ER C-terminal α-helix directly contributes to the toggle switch function. Indeed, the deletion the C-terminal α-Helix alone alleviates BKI1 inhibition and diminishes signal attenuation mediated by PUB30 and PUB31 as much as the removal of the entire ER C-terminal tail, thereby resulting in stronger signal output and hyperactivity in both stomatal development and inflorescence growth (Figs. 2, 6, S4). Does this mean that the region outside of the α-Helix plays an insignificant role? We argue that the remaining C-terminal tail provides a critical interaction interface with the key ER signaling components. Indeed, a removal of the C-terminal 43 amino acids (ER_ΔC43_) further compromised the association of ER with both BKI1 and PUB30/31 (Fig. 2). The additional ER C-terminal tail region outside of the α-Helix possesses multiple transphosphorylation sites important for the ER protein stability as well as protein-protein interactions (Figs. 4, 5, S9, S11, Table S1). Thus, this region likely forms an unstructured flexible linker to function as a phosphorylation-regulated hinge to change binding partners (Fig. 6). A similar mechanism has been proposed for human ribosomal S6 kinase1, in which C-terminal tail phosphorylations within the disordered domain trigger both charge- and conformational changes, exposing the protein-protein interaction surfaces (55). In such a case, the phosphosites may act as an on-off switch or a dimmer to collectively attenuate the signal output. Likewise, a mode of autoinhibition via the C-terminal tail and following release via phosphorylation also exists in H^+^-ATPase during blue light activation of stomatal opening, suggesting a broad role of CTs as roadblocks and removal of roadblocks via phosphorylation in plants as well (56).

Our study provides further insight into the role of C-terminal tail phosphoregulation in plant receptor kinases. The length and sequence of the C-terminal tails are highly variable amongst the plant LRR-RK families, with some members (e.g. PSKR1, CLV1, and BAM3) lacking or having very short C-terminal tails (Fig. S2). It is known that ER-family LRR-RKs simultaneously perceives multiple ligands emanating from the neighboring cells and tissues, some with redundant or antagonistic functions (24, 57). The unique structure and function of the ER C-terminal tail may be important for avoiding signal interference to achieve precise information processing during diverse developmental processes mediated by ER-family RKs. Further studies in ER-family and other RKs will elucidate the conserved and unique modes of receptor kinase activation and attenuation.

## Acknowledgements

We thank Libo Shan and Xiangzong Meng for MBP-BAK1 constructs; Krishna Mohan Sepuru for expert advice on BLI assays and CD spectrometry; Keiko Kuwata for expert assistance in mass-spectrometry; and Naoyuki Uchida, Krishna Mohan Sepuru, and Pengfei Bai for critiques on our manuscript. This work was initially supported by US National Science Foundation (IOS-0744892) and Gordon and Betty Moore Foundation (GBMF3035) to K.U.T., and NIH grant R35GM119536 to J.V. K.U.T. acknowledges the support from Howard Hughes Medical Institute and Johnson and Johnson Centennial Chair from The University of Texas at Austin.

## Author contributions

Conceived, K.U.T.; Designed experiments, L.C., M.M., J.A., K.U.T.; Generated plasmids and transgenic lines, L.C., M.M., J.A., K.U.T.; Characterized plant phenotypes and genotypes, L.C., M.M., A.C., K.U.T.; Biochemistry and biophysics, L.C., J.A.; *In vitro* phosphorylation assays, M.M., J.A.; *In vivo* phosphorylation assays, M.M., J.A., J.V., K.H., P.D., J.S., F.L.H.M.; Analyzed data, L.C., M.M., J.A., J.V., K.H., J.S., F.L.H.M., K.U.T.; Phylogenetic analysis, K.U.T.; Visualization, L.C., M.M., K.U.T.; Writing-Draft, L.C., K.U.T.; Project Administration, K.U.T.; Funding Acquisition, J.V., F.L.H.M., K.U.T.

## Materials and Methods

Plant materials and growth conditions, Plasmid construction and generation of transgenic plants, microscopy, quantitative analysis and statistics, expression, purification, and refolding of peptides, co-immunoprecipitation, protein gel electrophoresis, and immunoblots, *in vivo* ubiquitination assays, *in vitro* kinase assays and detection of phosphosites by mass spectrometry, detection of *in vivo* phosphorylation sites by mass spectrometry, biolayer interferometry (BLI), ER protein stability assay in protoplasts, RT-qPCR analysis, Circular Dichroism (CD) spectroscopy are described in *SI Materials and Methods*.

